## Supplementary Material for "Loss-of-function, gain-of-function and dominant-negative mutations have profoundly different effects on protein structure: implications for variant effect prediction"

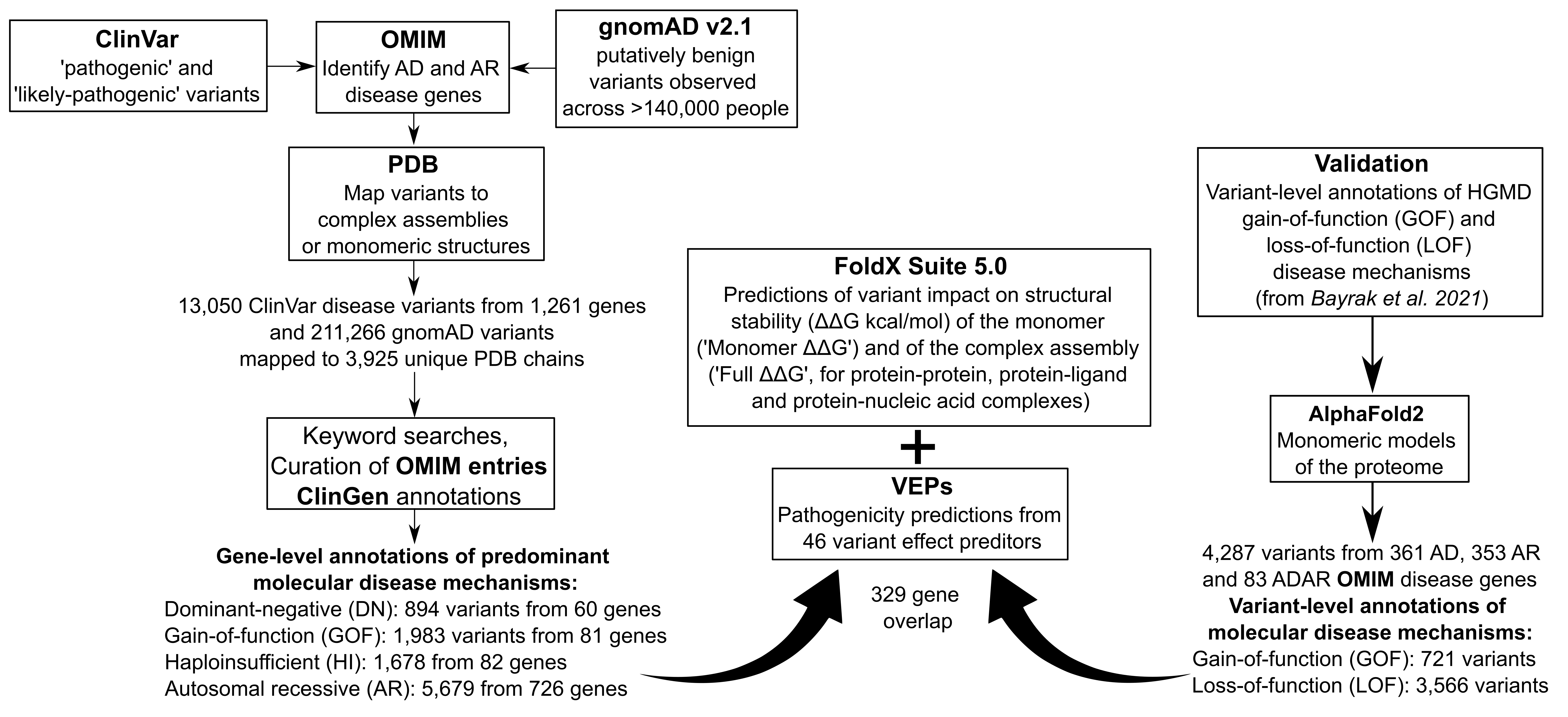
**Figure S1. Schematic representation of the data collection, gene annotation, prediction and validation setups.** See ‘Methods’.

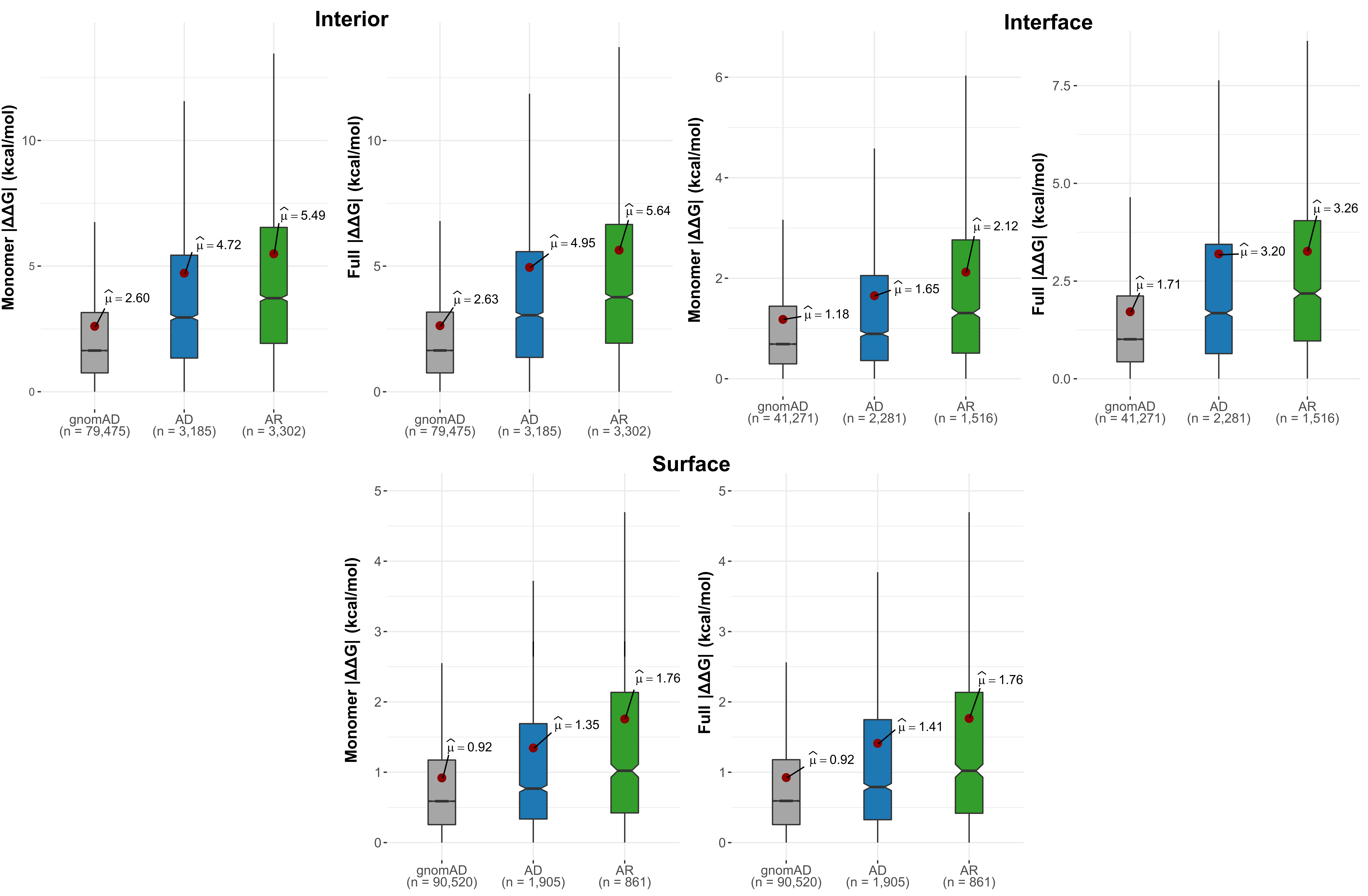
**Figure S2. Differences in |ΔΔG| values between gnomAD, autosomal dominant and autosomal recessive mutations are observed across interior, interface and surface locations.** All pairwise group comparisons showed significant differences (P < 5.0 x 10^-6^, Holm-corrected Dunn’s test).

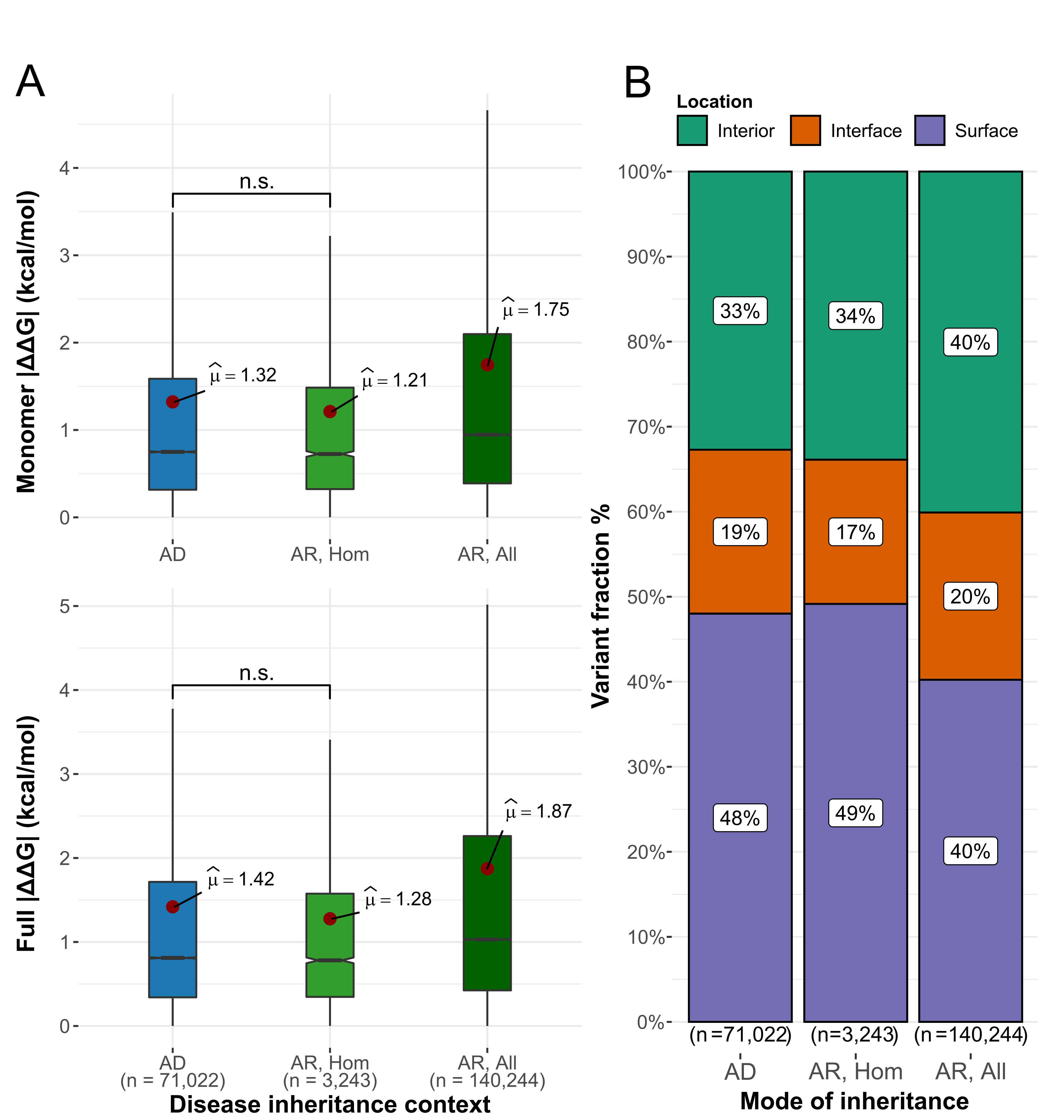

**Figure S3.** **Accounting for variant zygosity reveals highly similar |ΔΔG| values and location distributions for gnomAD variants in autosomal dominant and autosomal recessive disease genes.** **(A)** Stability perturbation differences between gnomAD variants of different inheritance contexts and zygosity. ‘AR, All’ considers all gnomAD variants from the AR genes, while ‘AR, Hom’ only includes those variants that have been observed in a homozygous state in gnomAD at least once. Pairwise group comparisons are significant unless specified (P < 1.4 x 10^-34^, Holm-corrected Dunn’s test). **(B)** Proportions of gnomAD variants throughout spatial structure locations for gene variants characterised by different inheritance context groups (all Chi-square comparisons are significant; Cramer’s V effect sizes are 0.01 and 0.08 for AD vs AR comparisons, using only homozygous recessive variants or all recessive variants, respectively).

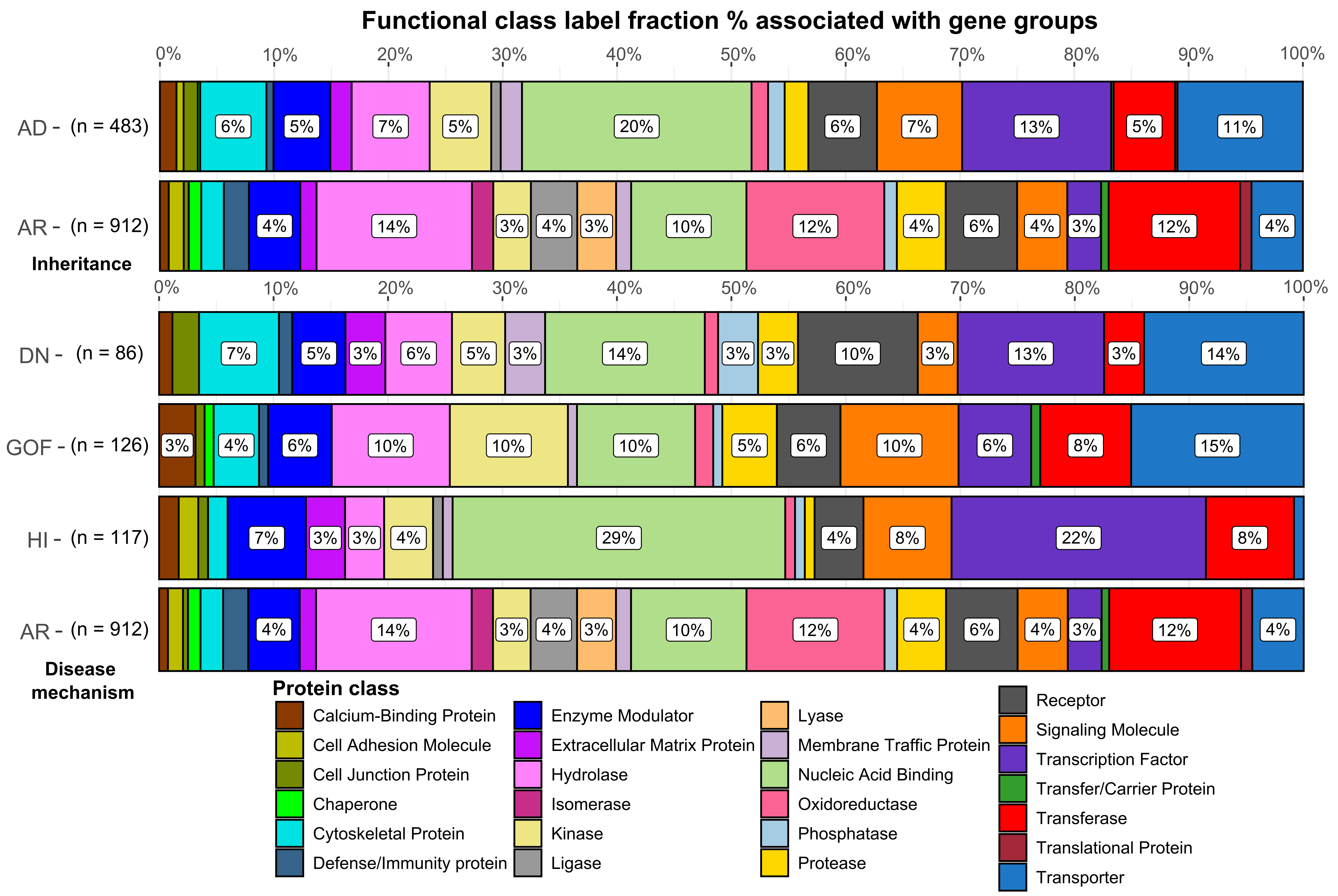
 **Figure S4. Inheritance and molecular mechanism gene groups are characterised by distinct functional protein class label prevalence.** Chi-square test comparisons were performed within the inheritance and mechanism groups, with only the non-LOF DN *vs* GOF mechanism genes showing insufficient difference (P = 0.147, highest significant P = 4.116 x 10^-3^ after Holm’s correction for multiple comparisons). Sample sizes denote functional class label number. The same gene can be associated with multiple functional classes.

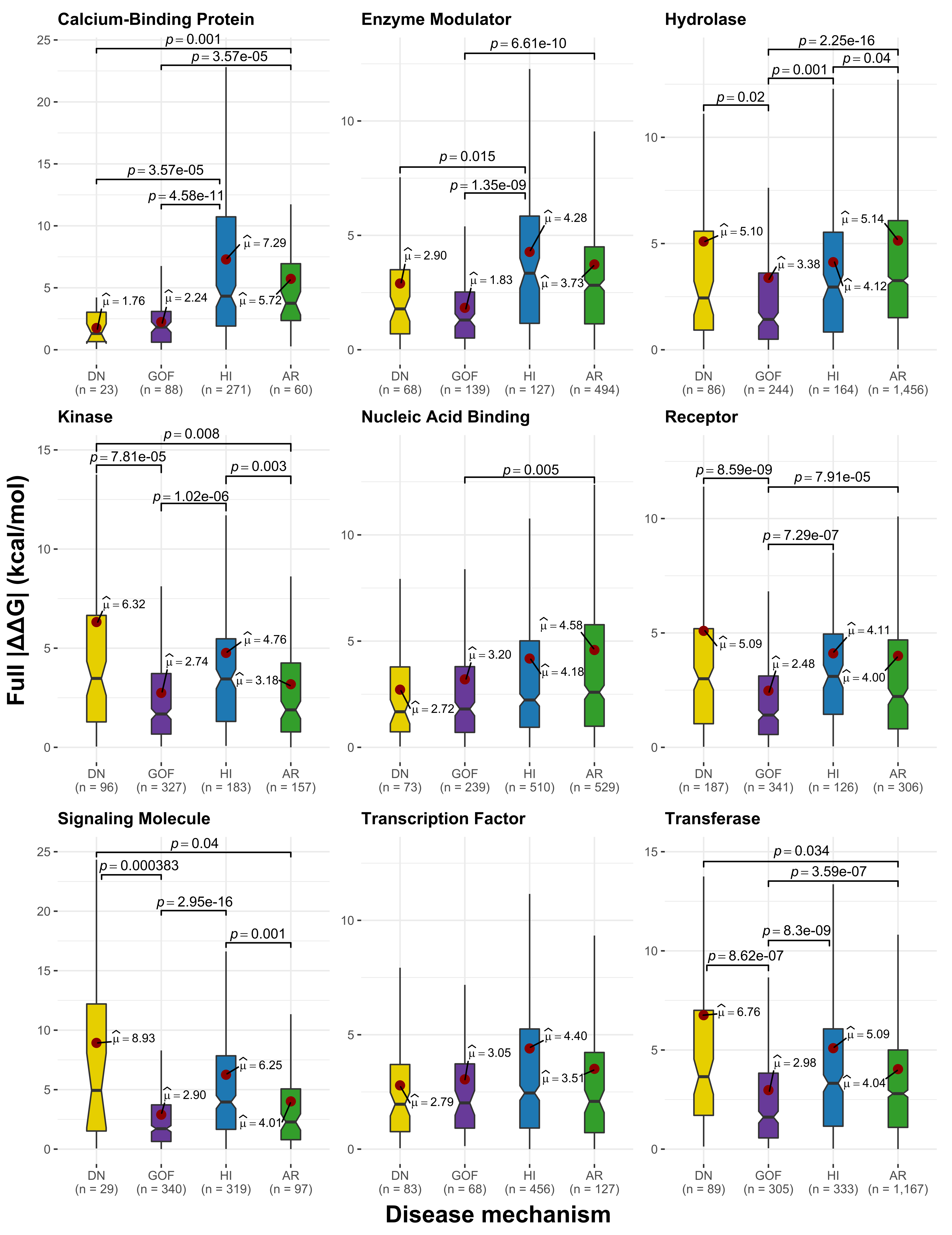
 **Figure S5. Underlying functional protein class does not drive the observed the variance differences in distinct molecular mechanism perturbation magnitude**. Only functional class groups with at least 20 variants in each molecular mechanism were analyzed. Statistically significant comparisons are shown (Holm-corrected Dunn’s test).

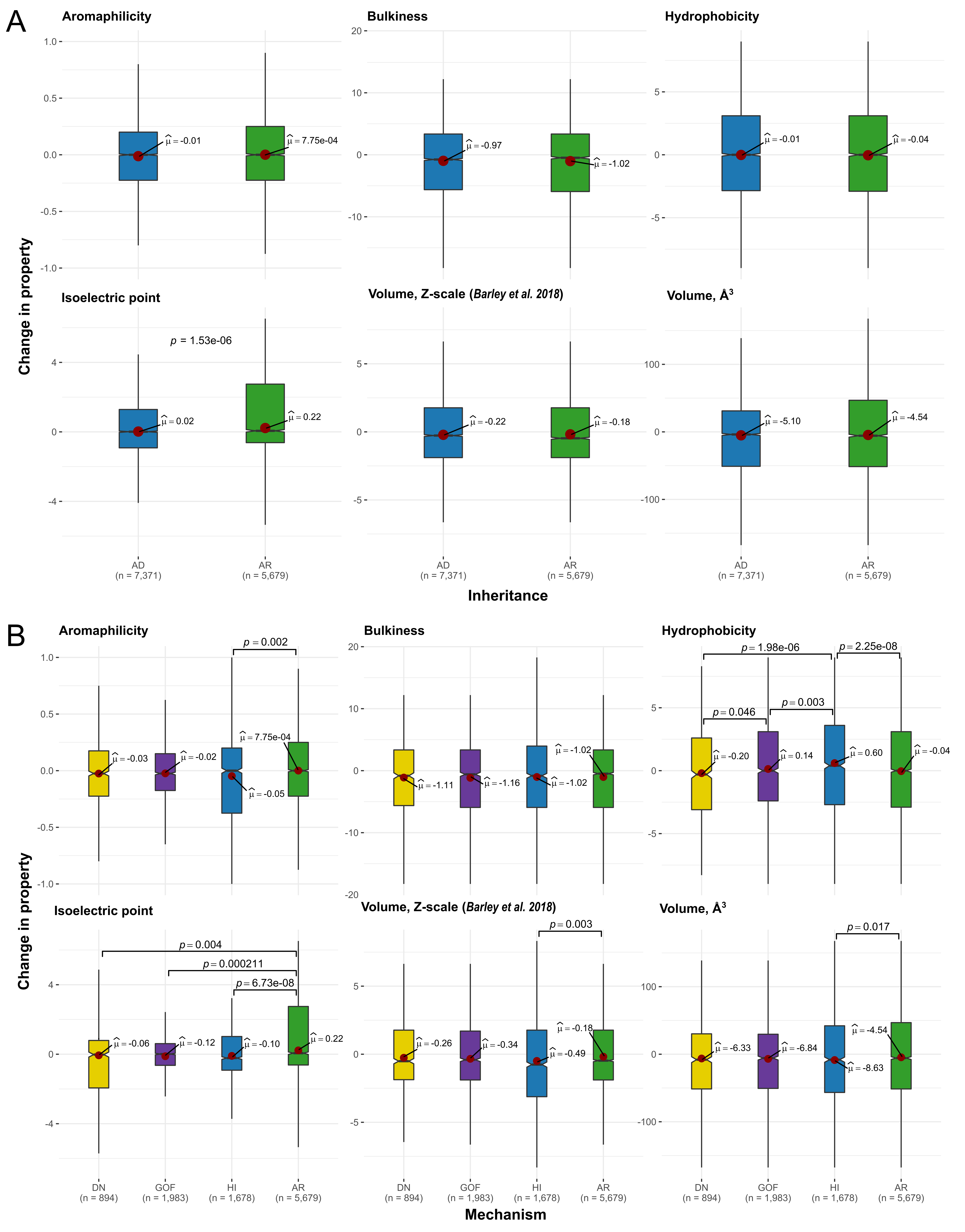
 **Figure S6. Physicochemical property index changes between wild-type and mutant residues in inheritance (A)** **and mechanism (B)** **groups.** Index sources and values for each amino acid residue can be found in the full dataset file (see ‘Data Availability’)**.** Negative property change values indicate an increase in property upon mutation, compared to the wild type residue. Statistically significant comparisons are shown (Holm-corrected Dunn’s test).

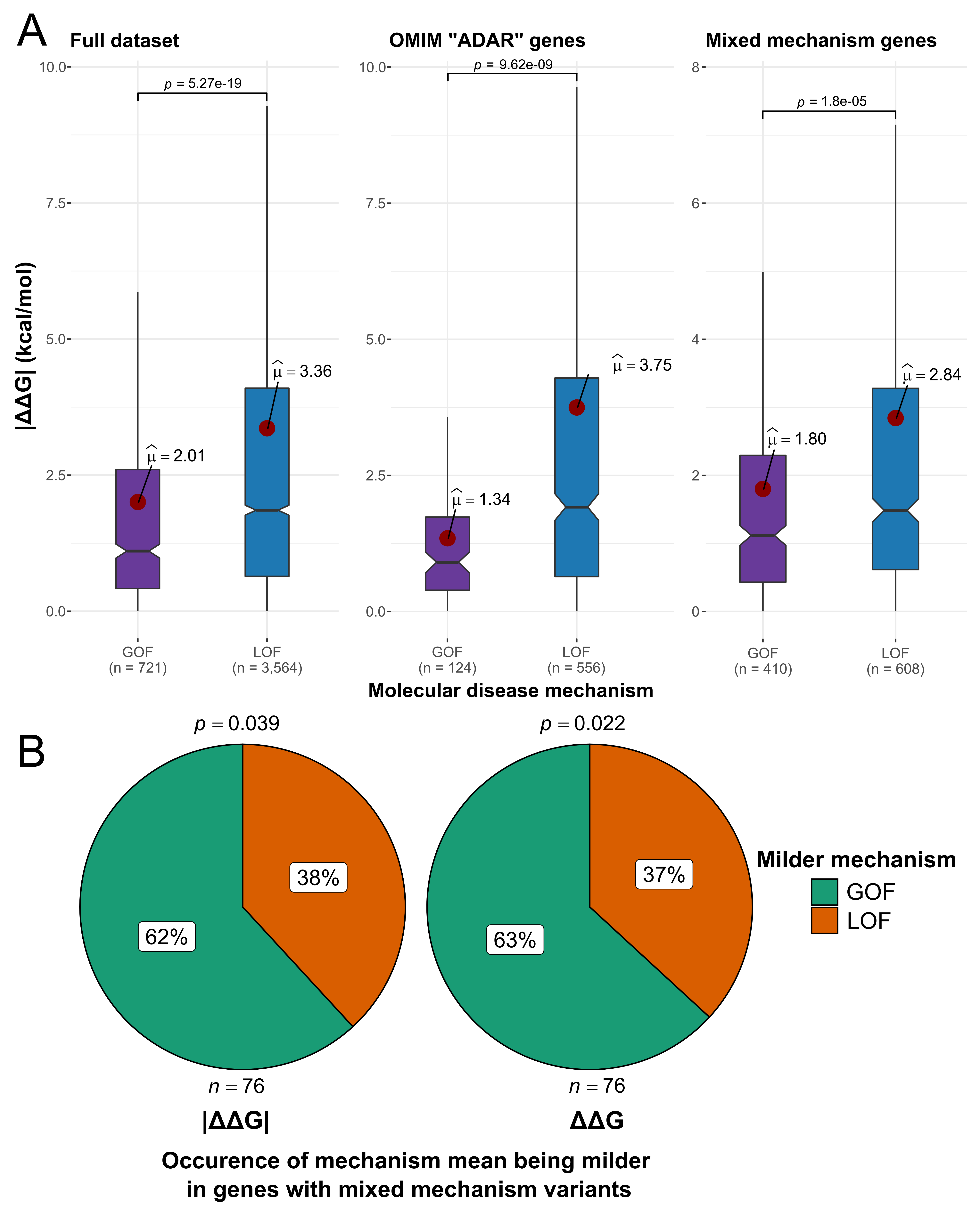
**Figure S7. Gain-of-function variants are also milder than loss-of-function mutations in mixed inheritance and mechanism genes.** The variant-level validation dataset is based on the HGMD GOF/LOF disease mechanism data from Bayrak et al., 2021^1^. **(A)** Comparison of predicted absolute ΔΔG value differences between GOF and LOF variants in several distinct contexts: the full dataset; only looking at mixed inheritance OMIM ‘ADAR’ genes; only genes with both GOF and LOF variants. Significant group comparisons are denoted (Wilcoxon rank-sum test). **(B)** Proportion of genes with both kind of variants (GOF & LOF), according to which mechanism group demonstrates a lower predicted ΔΔG mean within the same gene (Chi-square p-values depicted).

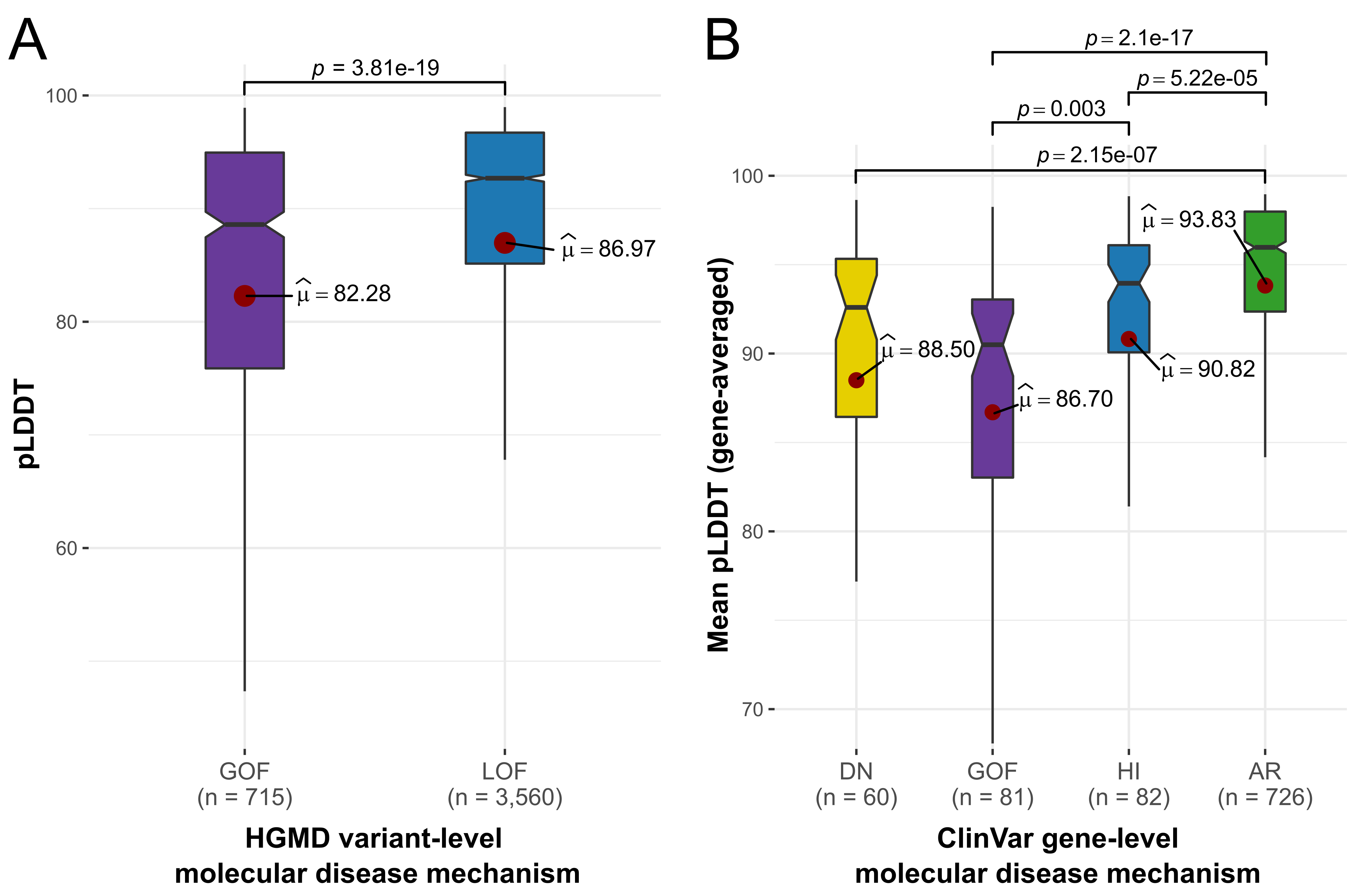
 **Figure S8. Gain-of-function variants occur in less structurally ordered positions than other mechanism mutations, according to the AlphaFold pLDDT modelling quality metric.** The AlphaFold pLDDT metric has been shown to be an accurate proxy for structural disorder^2^. Statistically significant comparisons are shown (Holm-corrected Dunn’s test). **(A)** pLDDT differences between HGMD GOF and LOF variants. Sample sizes represent variant number **(B)** pLDDT comparison across gene-level ClinVar mechanism groups. To control for gene-level annotation biases and uneven variant counts across genes, the pLDDT value is presented as a per-gene mean. Sample sizes represent gene number.

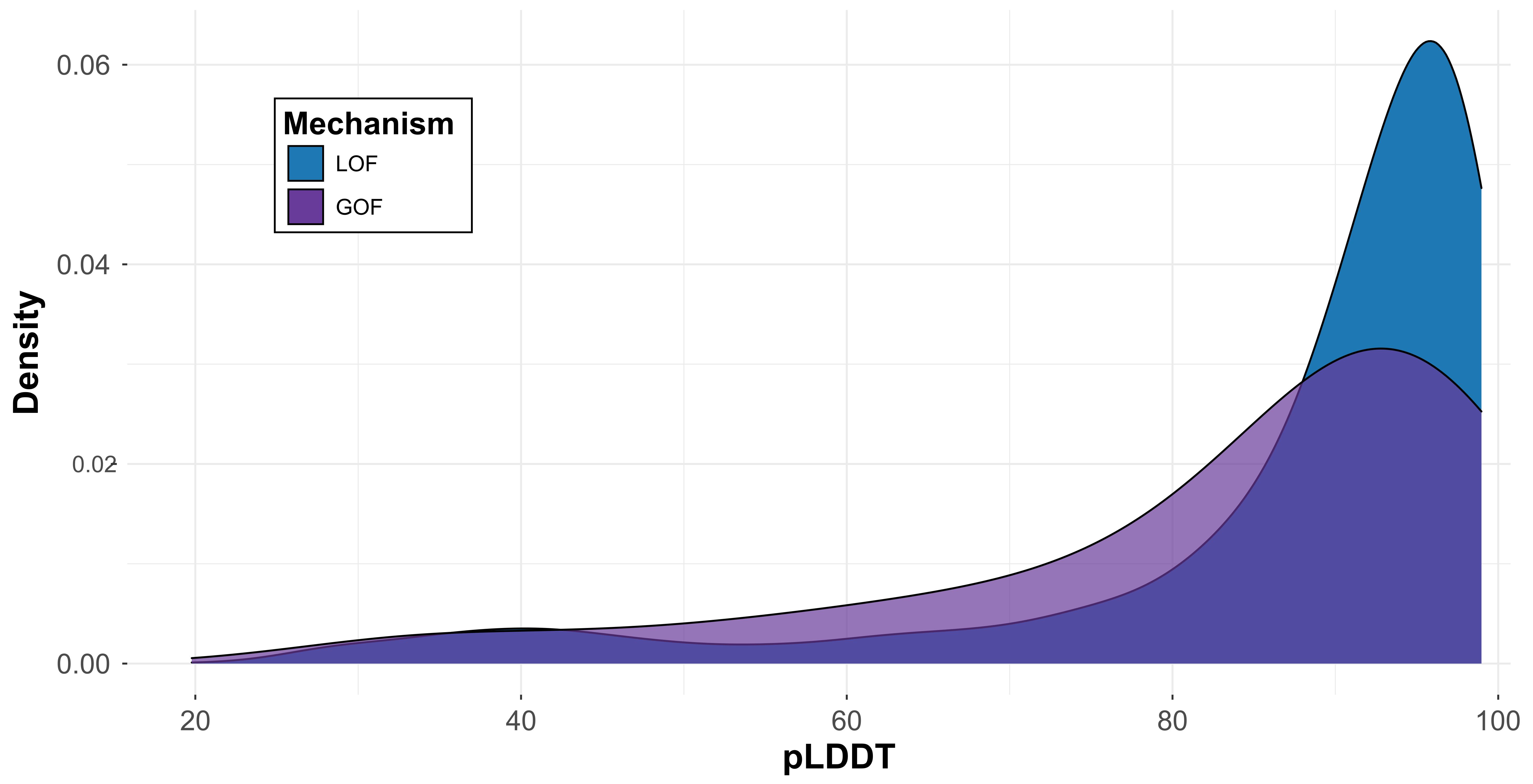
**Figure S9. Majority of disease variants occur in positions characterised by high pLDDT, independent of variant-level mechanism.** The values were derived using the variant-level HGMD dataset.

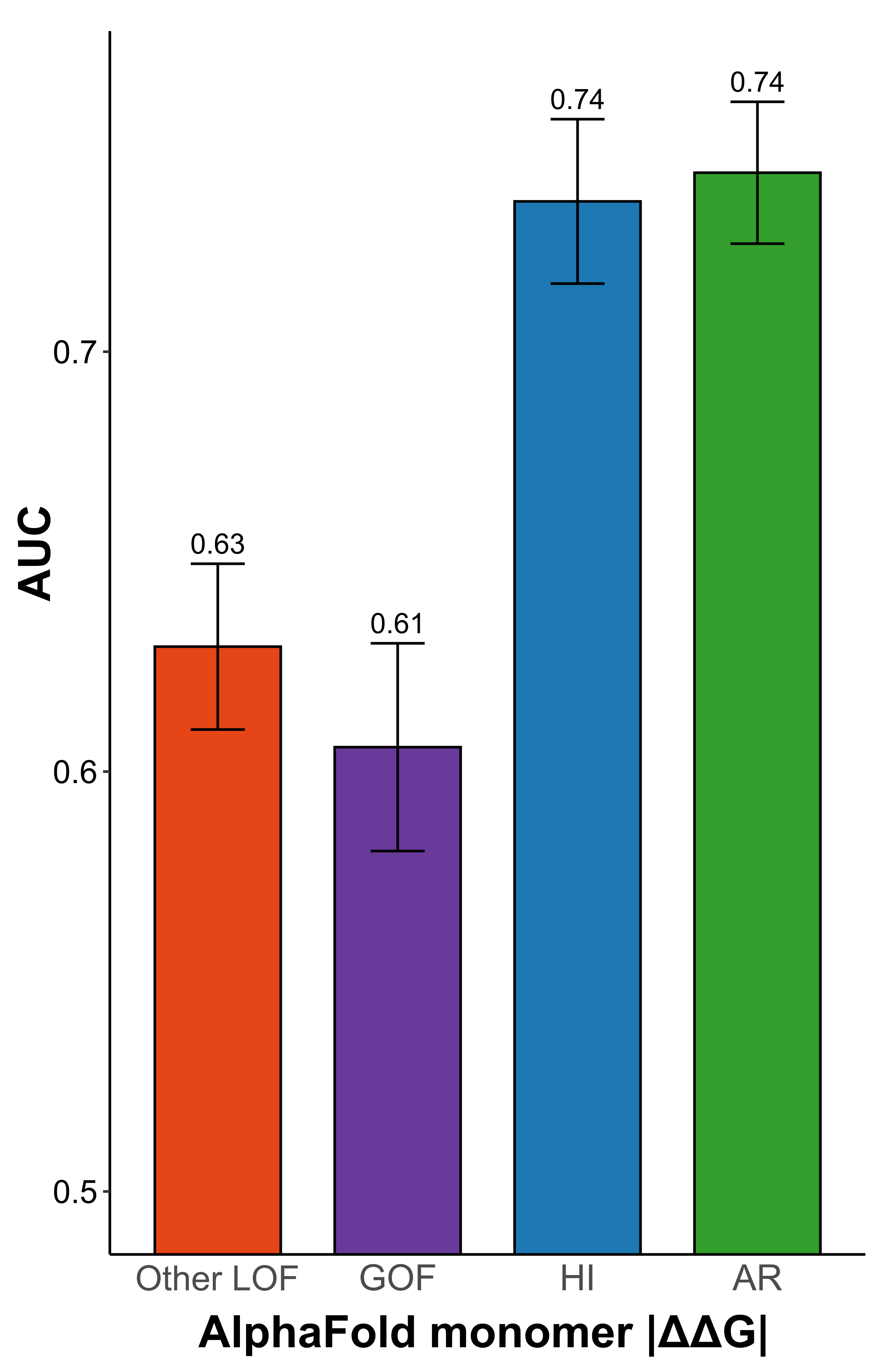

**Figure S10.** **Independent dataset recapitulates observed non-LOF mechanism structural mildness.** The variant-level validation dataset is based on the HGMD GOF/LOF disease mechanism data from Bayrak et al., 2021^1^. AUC values calculated from ROC curves for discriminating between different types of pathogenic HGMD mutations, and putatively benign gnomAD variants, based on predicted stability change score. Only homozygous gnomAD variants were included for the AR analysis. Error bars denote 95% confidence intervals, which were computed with 2000 stratified bootstrap replicates.

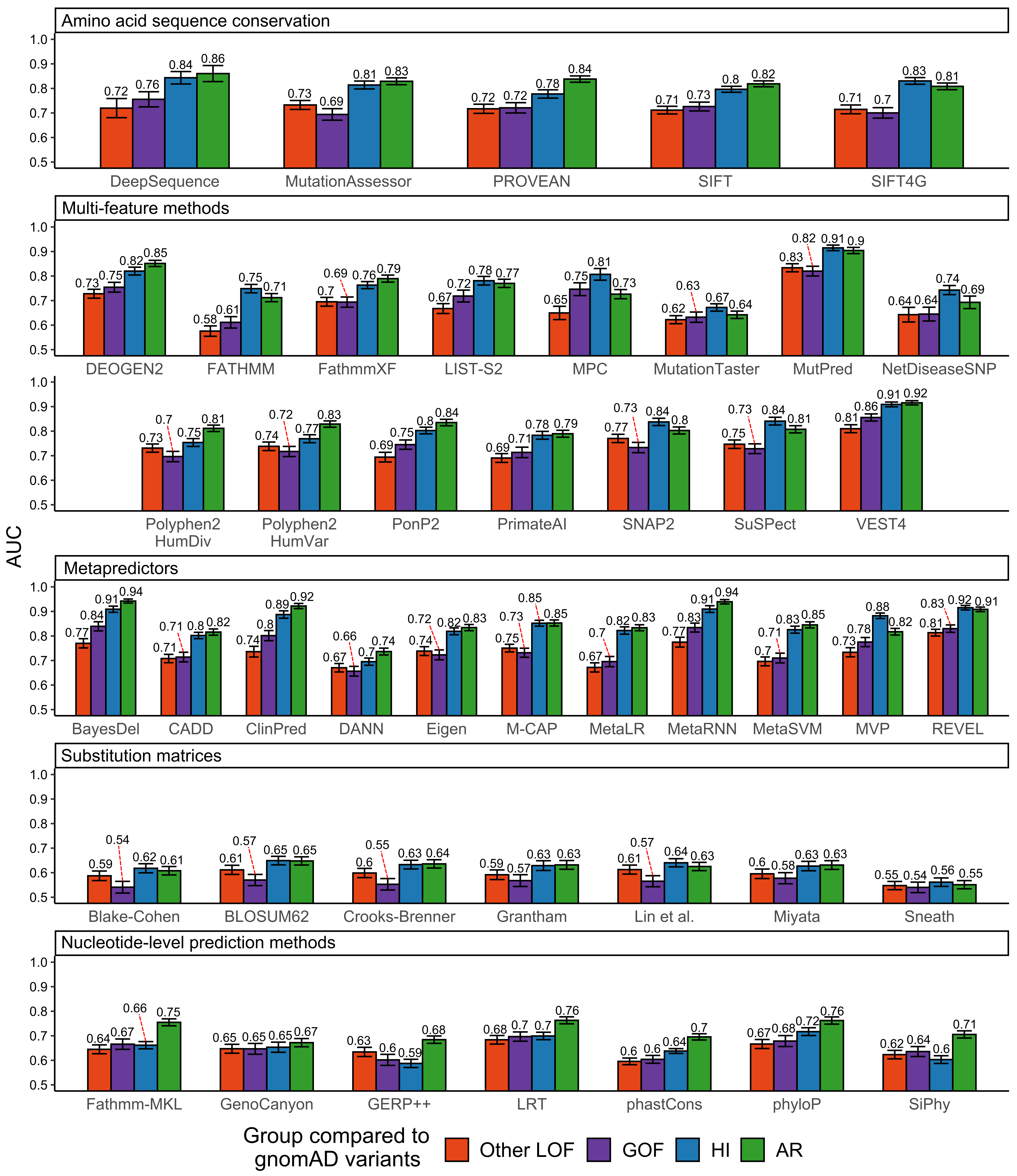
 **Figure S11. Hybrid variant- and gene-level disease mechanism classification approach validates previously observed VEP performance results.** The validation dataset is based on an external variant-level disease mechanism GOF/LOF label dataset from Bayrak et al. 2021^1^ (see ‘Methods’). AUC values calculated from ROC curves for discriminating between different types of pathogenic HGMD mutations and putatively benign gnomAD variants, using the outputs of different computational variant effect predictors. Only homozygous gnomAD variants were included for the AR analysis. Error bars denote 95% confidence intervals, which were computed with 2000 stratified bootstrap replicates.

**Table S1. Optimal FoldX |ΔΔG| thresholds to distinguish between pathogenic and putatively benign mutations for genes associated with different molecular disease mechanisms.**

| Molecular disease mechanism group | Predictor | Optimal threshold (kcal/mol) | Specificity (low est.) | Specificity (median) | Specificity (high est.) | Sensitivity (low est.) | Sensitivity (median) | Sensitivity (high est.) |
| --- | --- | --- | --- | --- | --- | --- | --- | --- |
| DN | FoldX Monomer \|ΔΔG\| | 1.03 | 52.01 | 55.15 | 58.39 | 60.46 | 61.78 | 63.06 |
| GOF | FoldX Monomer \|ΔΔG\| | 0.98 | 56.68 | 58.80 | 60.97 | 57.88 | 58.72 | 59.58 |
| HI | FoldX Monomer \|ΔΔG\| | 1.38 | 62.22 | 64.66 | 66.92 | 72.76 | 73.63 | 74.54 |
| AR (homozygous gnomAD) | FoldX Monomer \|ΔΔG\| | 1.41 | 64.61 | 65.87 | 67.14 | 72.12 | 73.64 | 75.21 |
| DN | FoldX Full \|ΔΔG\| | 1.28 | 57.05 | 60.18 | 63.53 | 64.70 | 65.95 | 67.29 |
| GOF | FoldX Full \|ΔΔG\| | 1.16 | 55.87 | 57.89 | 60.01 | 61.29 | 62.06 | 62.84 |
| HI | FoldX Full \|ΔΔG\| | 1.59 | 66.03 | 68.30 | 70.50 | 74.81 | 75.65 | 76.52 |
| AR (homozygous gnomAD) | FoldX Full \|ΔΔG\| | 1.55 | 67.11 | 68.30 | 69.50 | 73.27 | 74.68 | 76.19 |

**Table S2. Variant effect predictors used in this study.**

| Predictor | Feature-based classification | ClinVar disease dataset size | gnomAD dataset size | Source access | Reference |
| --- | --- | --- | --- | --- | --- |
| BayesDel | Metapredictor | 5017 | 39289 | dbNSFP database | (Feng, B. J., 2017)^3^ |
| Blake-Cohen | Substitution matrix | 13048 | 211134 | https://www.genome.jp/entry/aaindex:BLAJ010101 | (Blake & Cohen, 2001)^4^ |
| BLOSUM62 | Substitution matrix | 13048 | 211134 | https://www.genome.jp/entry/aaindex:HENS920102 | (Henikoff & Henikoff, 1992)^5^ |
| CADD | Metapredictor | 12806 | 208522 | https://cadd.gs.washington.edu/snv | (Kircher et al., 2014)^6^ |
| ClinPred | Metapredictor | 5017 | 39289 | dbNSFP database | (Alirezaie et al., 2018)^7^ |
| Crooks-Brenner | Substitution matrix | 13048 | 211134 | https://www.genome.jp/entry/aaindex:CROG050101 | (Crooks & Brenner, 2005)^8^ |
| DANN | Metapredictor | 12801 | 208138 | dbNSFP database | (Quang et al., 2015)^9^ |
| DeepSequence | Amino acid sequence conservation | 4613 | 19029 | https://github.com/debbiemarkslab/DeepSequence | (Riesselman et al., 2018)^10^ |
| DEOGEN2 | Multi-feature | 12095 | 202031 | https://deogen2.mutaframe.com/ | (Raimondi et al., 2017)^11^ |
| Eigen | Metapredictor | 12772 | 207420 | dbNSFP database | (Ionita-Laza et al., 2016)^12^ |
| FATHMM | Multi-feature | 12113 | 196893 | http://fathmm.biocompute.org.uk/inherited.html | (Shihab et al., 2013)^13^ |
| Fathmm-MKL | Multi-feature | 12801 | 208138 | dbNSFP database | (Shihab et al., 2015)^14^ |
| FathmmXF | Nucleotide-level prediction method | 12712 | 207661 | http://fathmm.biocompute.org.uk/fathmm-xf/ | (Rogers et al., 2018)^15^ |
| GenoCanyon | Nucleotide-level prediction method | 12801 | 208138 | dbNSFP database | (Lu et al., 2015)^16^ |
| GERP++ | Nucleotide-level prediction method | 12801 | 208129 | dbNSFP database | (Davydov et al., 2010)^17^ |
| Grantham | Substitution matrix | 13048 | 211134 | https://www.genome.jp/entry/aaindex:GRAR740104 | (Grantham, 1974)^18^ |
| Lin et al. | Substitution matrix | 13048 | 211134 | https://www.genome.jp/entry/aaindex:LINK010101 | (Lin et al., 2001)^19^ |
| LIST-S2 | Metapredictor | 4407 | 34468 | dbNSFP database | (Malhis et al., 2020)^20^ |
| LRT | Nucleotide-level prediction method | 12546 | 201271 | dbNSFP database | (Chun & Fay, 2009)^21^ |
| M-CAP | Metapredictor | 12746 | 206078 | http://bejerano.stanford.edu/mcap/ | (Jagadeesh et al., 2016)^22^ |
| MetaLR | Metapredictor | 12772 | 207420 | dbNSFP database | (Dong et al., 2015)^23^ |
| MetaRNN | Metapredictor | 5017 | 39289 | dbNSFP database | (Li et al., 2021)^24^ |
| MetaSVM | Metapredictor | 12772 | 207420 | dbNSFP database | (Dong et al., 2015)^23^ |
| Miyata | Substitution matrix | 13048 | 211134 | https://www.genome.jp/entry/aaindex:MIYT790101 | (Miyata et al., 1979)^25^ |
| MPC | Multi-feature | 9088 | 162846 | dbNSFP database | (Samocha et al., 2017)^26^ |
| MutationAssessor | Amino acid sequence conservation | 11873 | 195899 | dbNSFP database | (Reva et al., 2011)^27^ |
| MutationTaster | Multi-feature | 12761 | 206319 | dbNSFP database | (Schwarz et al., 2014)^28^ |
| MutPred | Multi-feature | 11554 | 170575 | dbNSFP database | (Pejaver et al., 2020)^29^ |
| MVP | Metapredictor | 12528 | 205851 | dbNSFP database | (Qi et al., 2018)^30^ |
| NetDiseaseSNP | Multi-feature | 12789 | 205379 | http://www.cbs.dtu.dk/services/NetDiseaseSNP/ | (Johansen et al., 2013)^31^ |
| phastCons | Nucleotide-level prediction method | 12806 | 208522 | dbNSFP database | (Siepel et al., 2005)^32^ |
| phyloP | Nucleotide-level prediction method | 12806 | 208522 | http://papi.unipv.it/ | (Pollard et al., 2010)^33^ |
| PolyPhen2 HumDiv | Multi-feature | 11771 | 198850 | http://genetics.bwh.harvard.edu/pph2/ | (Adzhubei et al., 2010)^34^ |
| PolyPhen2 HumVar | Multi-feature | 11771 | 198850 | http://genetics.bwh.harvard.edu/pph2/ | (Adzhubei et al., 2010)^34^ |
| PonP2 | Multi-feature | 12134 | 196040 | http://structure.bmc.lu.se/PON-P2/ | (Niroula et al., 2015)^35^ |
| PrimateAI | Multi-feature | 12638 | 206477 | dbNSFP database | (Sundaram et al., 2018)^36^ |
| PROVEAN | Amino acid sequence conservation | 12044 | 197456 | http://provean.jcvi.org/index.php | (Choi et al., 2012)^37^ |
| REVEL | Metapredictor | 12772 | 207420 | https://sites.google.com/site/revelgenomics/ | (Ioannidis et al., 2016)^38^ |
| SIFT | Amino acid sequence conservation | 12851 | 208985 | https://sift.bii.a-star.edu.sg/www/code.html | (Sim et al., 2012)^39^ |
| SIFT4G | Amino acid sequence conservation | 12418 | 203379 | dbNSFP database | (Vaser et al., 2016)^40^ |
| SiPhy | Nucleotide-level prediction method | 12799 | 208117 | dbNSFP | (Garber et al., 2009)^41^ |
| SNAP2 | Multi-feature | 13009 | 210582 | https://www.rostlab.org/services/snap/ | (Hecht et al., 2015)^42^ |
| Sneath | Substitution matrix | 13048 | 211134 | (Sneath, 1966) | (Sneath, 1966)^43^ |
| SuSPect | Multi-feature | 13007 | 210532 | http://www.sbg.bio.ic.ac.uk/suspect/about.html | (Yates et al., 2014)^44^ |
| VEST4 | Multi-feature | 12633 | 206645 | https://www.cravat.us/CRAVAT/ | (Carter et al., 2013)^45^ |
